# Polar Geometry of Time-Odd EEG Dynamics

**DOI:** 10.64898/2026.09.08.749160

**Authors:** Dmitry Goldstein

**Affiliations:** Holon Institute of Technology, Holon, Israel

## Abstract

**Objective:** Longitudinal EEG recordings vary across sessions because of electrode reapplication, referencing, recording conditions, and physiological state. We asked whether multichannel EEG contains a time-odd geometric structure that remains subject-discriminative across repeated recordings and extended inter-session intervals.

**Approach:** We constructed a local state–derivative operator *T*_*ij*_ = corr(*m*_*i*_, Δ*x*_*j*_) and isolated its exact skew component, 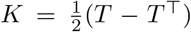. Polar decomposition, *K* = *Q*_*K*_*M*_*K*_, separated interaction magnitude from normalized orientation. We applied the same decomposition to signed imaginary coherency, Ω = Im(*G*), providing a frequency-domain test based on a separate estimation procedure. We evaluated cross-session subject matching in four public longitudinal EEG datasets, including the Dortmund cohort of 206 participants with recordings separated by approximately five years. Alpha–beta fusion used fixed equal weights without calibration or learned weighting.

**Main results:** The empirical operator *T* was already strongly dominated by its skew component, indicating that *K* isolates rather than creates the observed time-odd structure. Polar orientation improved longitudinal discriminability for *K* → *Q*_*K*_ in every principal comparison. For the approximately one-month RestCog interval, area under the receiver-operating-characteristic curve (AUC) increased from 0.9222 to 0.9597 and rank-1 identification (CMC@1) from 0.6333 to 0.8167. In Dortmund, AUC increased from 0.8901 to 0.9625 and CMC@1 from 0.4757 to 0.6214. Fixed alpha–beta fusion further increased Dortmund *Q*_*K*_ performance to CMC@1 = 0.8350, AUC = 0.9855, and equal-error rate (EER) = 5.90%. The spectral operator reproduced the raw-to-polar AUC advantage for Ω → *Q*_Ω_ in both alpha and beta across all four datasets. Matched-metric controls preserved the polar AUC advantage in all 40 raw-to-polar comparisons.

**Significance:** Across two mathematically linked skew EEG operators obtained using different estimation procedures, removing interaction-magnitude weighting while retaining normalized orientation improved cross-session discriminability. The gain reflects improved genuine–impostor geometry rather than necessarily greater absolute similarity between repeated recordings. Polar orientation provides a compact longitudinal descriptor requiring no performance-tuned internal parameter once preprocessing, temporal support, and frequency band are fixed.

## 1 Introduction

A central challenge in longitudinal EEG analysis is to identify multichannel features that remain reproducible across recording sessions. Even when the nominal acquisition protocol and montage are unchanged, repeated recordings can differ because of electrode reapplication, contact conditions, referencing, signal amplitude, vigilance, and other session-specific physiological and technical factors. Longitudinal stability and permanence are therefore central considerations for subject-specific EEG representations [1–3].

We ask whether time-asymmetric multichannel dynamics contain a reproducible geometric structure. Our working hypothesis is that interaction magnitudes may be particularly sensitive to session-specific conditions, whereas aspects of their geometric organization may be more stable. We therefore examine the orientation of time-odd multichannel interactions rather than interaction magnitude alone.

Interactions between signal level and temporal derivative have previously been considered in differential-covariance descriptions of multivariate neural dynamics [4, 5]. Following this framework, we define midpoint and first-difference variables separated by the sampling interval Δ*t*,

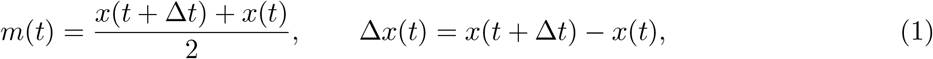

and form the empirical state–derivative operator

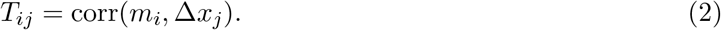

There is a direct theoretical reason to expect this operator to be predominantly time-odd. Under continuous-time stationarity, the state–derivative cross-moment

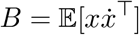

satisfies *B*^*T*^ = −*B*. Accordingly, we isolate the exact skew component

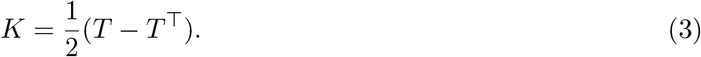

This operation projects *T* onto the theoretically expected time-odd component rather than constructing an unrelated representation. Whether the empirical correlation-normalized operator already exhibits this near-skew structure is empirically testable.

The central geometric object in this work is the polar orientation of *K*. In an orthogonal basis, a real skew-symmetric matrix decomposes into two-dimensional active planes with canonical blocks *σ*_*j*_*J*, where *σ*_*j*_ *>* 0 weights the corresponding skew plane. The matrix polar decomposition [6] separates normalized orientation from this magnitude weighting:

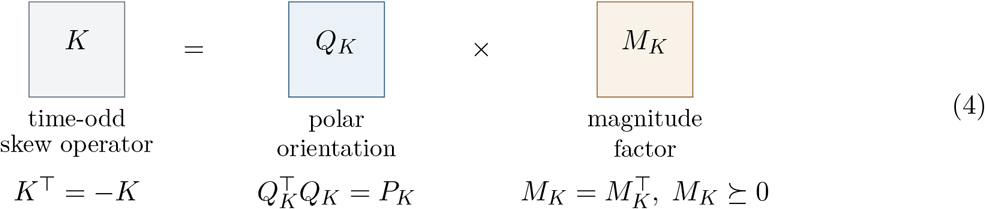

Here *P*_*K*_ denotes the orthogonal projector onto the active support of *K*. On each active skew plane, polar normalization maps *σ*_*j*_*J* to *J*. The factor *Q*_*K*_ therefore retains normalized orientation geometry, whereas *M*_*K*_ contains the corresponding magnitude information. Once preprocessing, temporal support, and frequency band are fixed, this separation introduces no performance-tuned internal parameter.

The state–derivative construction also admits a direct spectral interpretation. For scalar processes *u* and *v*, let *S*_*u,v*_(*f*) denote their cross-spectral density, and write 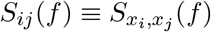. For the midpoint and first-difference variables,

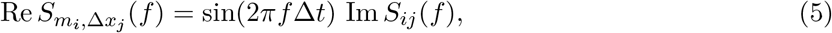

up to the Fourier-sign convention, with sin(2*πf* Δ*t*) ≈ 2*πf* Δ*t* for small Δ*t*. The local time-odd operator and signed imaginary spectral interaction therefore arise from the same antisymmetric cross-spectral structure while differing in frequency weighting and normalization.

Motivated by this relation, we introduce a frequency-domain skew operator estimated directly from signed imaginary coherency [7],

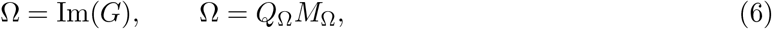

where *G* is the complex coherency matrix. The two skew operators are mathematically linked but not equivalent. The operator *K* represents a correlation-normalized state–derivative interaction with frequency weighting induced by temporal differencing, whereas Ω represents normalized signed imaginary spectral interaction estimated directly in the frequency domain. Their comparison tests whether the polar-orientation principle persists under different weighting, normalization, and estimation procedures.

Previous work has used cross-session EEG identification to assess the permanence and subject specificity of spectral and functional-connectivity representations [1–3, 8]. More broadly, connectivity-based identification has served as an assay of stable individual organization in neuroimaging [9]. We use subject matching here as a quantitative assay of longitudinal discriminability rather than as the primary application. If a representation preserves subject-specific organization, genuine enrollment– probe comparisons should remain better separated from impostor comparisons even when absolute within-subject similarity does not increase.

We evaluate these hypotheses across four public longitudinal EEG datasets spanning different cohort sizes and inter-session intervals. Zero-lag Pearson correlation provides a symmetric reference, while alpha and beta bands assess frequency specificity and fixed cross-band complementarity. Matched-metric, finite-lag, derivative-parity, derivative-discretization, magnitude-factor, and artificial-skew controls examine alternative explanations of the raw-to-polar effect. The analysis addresses three linked questions: whether the empirical state–derivative operator contains the predicted time-odd structure, whether polar orientation improves its longitudinal discriminability, and whether the same principle is reproduced by the linked spectral operator Ω.

## 2 Methods

### 2.1 Datasets and cross-session protocols

Four longitudinal EEG datasets were analyzed using prespecified enrollment–probe pairs, participant sets, channel orders, and temporal supports. The cohorts span intervals from short-term repeat recordings to an approximately five-year follow-up (Table 1).

**Table 1:** Characteristics of the four longitudinal EEG datasets and the principal prespecified enrollment– probe protocols. RestCog contributed two longitudinal comparisons but constitutes a single dataset.

| Dataset | OpenNeuro ID | $N$ | Ch. | Enrollment → probe | Interval | Support |
| --- | --- | --- | --- | --- | --- | --- |
| RestCog | ds004148 | 60 | 59 | ses01 EC → ses02 EC | short | $6 \times 50$ s |
| RestCog | ds004148 | 60 | 59 | ses01 EC → ses03 EC | $\sim 1$ month | $6 \times 50$ s |
| ds007176 | ds007176 | 27 | 58 | V0 CE → V3 CE | longitudinal | central 200 s;<br>$4 \times 50$ s |
| SRM | ds003775 | 42 | 64 | t1 EC → t2 EC | $\sim 2$–3<br>months | central 200 s;<br>$4 \times 50$ s |
| Dortmund | ds005385 | 206 | 64 | ses-1 EC → ses-2 EC | $\sim 5$ years | $3 \times 50$ s |

*Notes.* EC, eyes closed; EO, eyes open; CE, closed-eyes condition in ds007176. All principal analyses used signals resampled to 128 Hz. Imaginary coherency was estimated in fixed, non-overlapping 50-s windows using common multitaper parameters across cohorts.

#### RestCog (OpenNeuro ds004148)

The dataset and its public release are described in [10, 11]. Sixty participants with 59 common EEG channels contributed paired eyes-closed recordings. Session 01 was compared separately with sessions 02 and 03; session 03 was recorded approximately one month later, with electrode reapplication between visits. Six 50-s windows were analyzed per recording. A separate ses01 → ses03 eyes-open spectral branch retained 58 participants because two EO recordings were unreadable.

#### ds007176 (OpenNeuro)

The cohort and its public release are described in [12, 13]. The V0 CE and V3 CE recordings of 27 participants were compared. Horizontal and vertical electrooculography (EOG) channels were excluded from the native 60-channel recordings, leaving 58 EEG channels. The central 200 s were analyzed as four 50-s windows.

#### SRM (OpenNeuro ds003775 / NEMAR on003775)

The dataset and its public release are described in [14, 15]. Forty-two participants with 64 common BioSemi EEG channels contributed paired eyes-closed recordings at t1 and t2, separated by approximately 2–3 months. The central 200 s were analyzed as four 50-s windows.

#### Dortmund Vital Study (OpenNeuro ds005385)

The longitudinal dataset and its public release are described in [16, 17]. The prespecified comparison used ses-1 and ses-2 EyesClosed acq-pre recordings acquired approximately five years apart. The final cohort contained 206 participants and 64 channels. Sub-230 was excluded because of insufficient ses-1 duration, and sub-425 because of severe saturation within the prespecified support. Three 50-s windows were analyzed per recording.

### 2.2 Preprocessing and recording-level aggregation

All principal analyses used a sampling rate of *f*_*s*_ = 128 Hz. The principal frequency band was alpha (8–13 Hz), with beta (13–30 Hz) used for the cross-band extension. Time-domain operators were computed after fixed, zero-phase, fourth-order Butterworth band-pass filtering. The spectral branch instead applied band restriction directly to the multitaper estimates.

Window-level matrices *A*_*w*_ were averaged to obtain one recording-level operator,

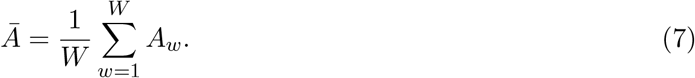

For the local state–derivative branch, antisymmetrization and polar factorization were applied after recording-level aggregation. Frequency bands, temporal supports, filter settings, and window counts were fixed independently of biometric performance; no recording-specific rank or support length was selected.

### 2.3 Cross-session matching and biometric evaluation

Each comparison produced an *N* × *N* score matrix

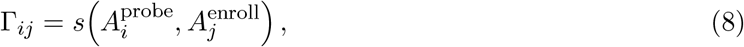

where *s*(*A, B*) denotes the representation-specific scalar pairwise comparison function defined below. Genuine scores lie on the diagonal and impostor scores off the diagonal. No classifier, identity-specific template learning, metric learning, or probe-dependent adaptation was used.

The normalized Frobenius cosine was

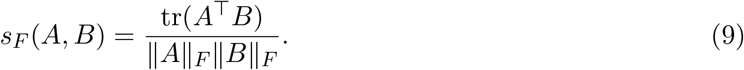

The directed operator *T* was matched by Pearson correlation of its directed off-diagonal entries. Exact-skew operators *K* and Ω were matched by Pearson correlation of their signed strict-upper-triangle entries. Positive polar factors were matched by Pearson correlation of the upper triangle, including the diagonal, whereas the orientation factors *Q*_*T*_, *Q*_*K*_, and *Q*_Ω_ were matched by full-matrix Frobenius cosine.

For distance-based reference representations, smaller values indicated a stronger match. Their signs were reversed for ranking and receiver-operating-characteristic evaluation so that all score matrices followed the common convention that larger values indicate greater similarity.

The matched-metric robustness analysis compared raw and polar operators using the same matcher before and after normalization: either full-matrix Frobenius cosine or strict-upper-triangle Pearson correlation. Identification performance was summarized by the cumulative match characteristic at ranks 1 and 5 (CMC@1 and CMC@5) and by mean genuine rank. Verification performance was summarized by the area under the receiver operating characteristic curve (AUC); equal-error rate (EER) was additionally reported for the cross-band and matched-metric analyses using a single global threshold per score matrix.

Alpha–beta fusion was fixed a priori as

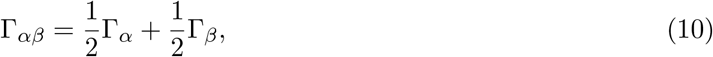

without score normalization, calibration, learned weights, band search, or dataset-specific optimization. Where paired inference was prespecified, AUC differences were evaluated using 10,000 probe-row bootstrap resamples, CMC differences using exact McNemar tests, and mean-rank differences using paired Wilcoxon signed-rank tests. Holm correction was applied across fusion-versus-alpha and fusion-versus-beta comparisons. Other cross-representation and cross-dataset comparisons were descriptive.

### 2.4 Zero-lag reference geometry

Each band-limited segment *X* ∈ ℝ^*p×n*^ was centered and standardized channel-wise and represented by

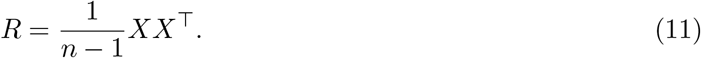

Matrices were compared using Frobenius distance, *d*_*F*_ (*R*_1_, *R*_2_) = ‖*R*_1_ *R*_2_‖ _*F*_, and the affine-invariant Riemannian metric (AIRM) [18],

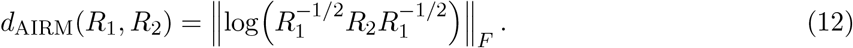

AIRM was evaluated after the fixed shrinkage *R*_*λ*_ = (1 − *λ*)*R* + *λI*, with *λ* = 0.01. This transformation multiplies Frobenius differences by the common factor 1 − *λ* and therefore does not alter their ranking.

### 2.5 Local state–derivative operator

For consecutive samples separated by Δ*t*,

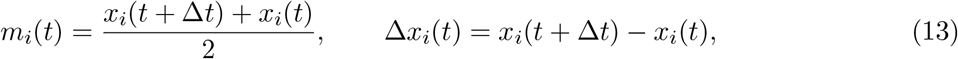

and, after independent standardization of the midpoint and difference sequences,

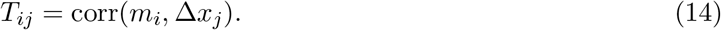

The factor 1*/*Δ*t* was omitted because positive rescaling does not affect Pearson correlation. This construction is related to differential-covariance and derivative-based descriptions of multivariate interactions [4, 5, 19].

Under continuous-time stationarity, *B* = E[*x*(*t*)*ẋ*(*t*)^*T*^] satisfies

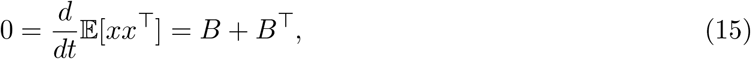

and hence *B*^*T*^ = *B*. The empirical correlation-normalized operator *T* need not be exactly skew-symmetric because the midpoint and difference sequences are normalized separately. Its exact time-odd

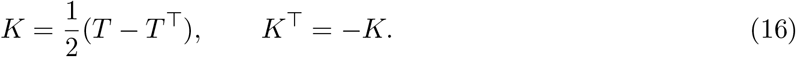

Using *S*_*u,v*_(*f*) to denote the cross-spectral density between scalar processes *u* and *v*, with 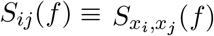, the exact frequency-domain relation for the midpoint–difference variables is

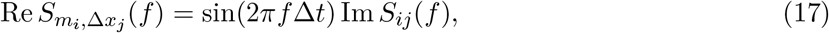

up to the Fourier-sign convention. Because sin(2*πf* Δ*t*) ≈ 2*πf* Δ*t* for small Δ*t, K* can be interpreted as a normalized time-domain realization of frequency-weighted imaginary cross-spectral coupling.

As a targeted mechanistic bridge, we also constructed a normalized spectral derivative operator,

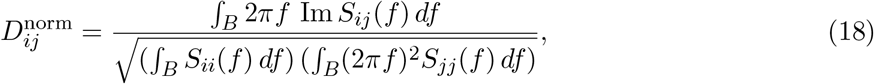

followed by antisymmetrization.

### 2.6 Polar geometry of skew operators

For a real skew-symmetric matrix *A* of rank 2*r*, an orthogonal basis exists in which

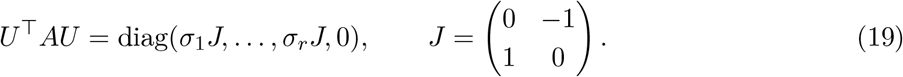

Its polar decomposition [6] is

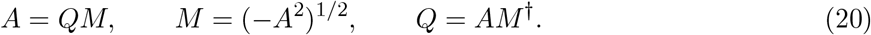

On each active skew plane, polar normalization maps *σ*_*j*_*J* to *J* : *M* retains plane magnitude, whereas *Q* removes the scalar weighting and retains normalized orientation.

This normalization is not guaranteed to improve longitudinal matching. When an active skew singular value approaches zero, its orientation becomes poorly conditioned. Moreover, a genuine pair can become less absolutely aligned after magnitude equalization while remaining better separated from impostors.

For numerical stability, polar factors were computed from the Hermitian matrix *H*_*A*_ = *iA*. If *H*_*A*_ = *V* Λ*V* ^*†*^, then, on the numerical active support,

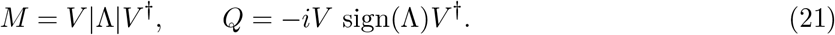

No performance-selected low-rank truncation was used. Odd-dimensional skew matrices were evaluated on their maximal numerical active support. For comparison, the ordinary polar decomposition *T* = *Q*_*T*_ *M*_*T*_ was also evaluated using the Moore–Penrose construction when required.

### 2.7 Frequency-domain skew operator: imaginary coherency

Signed imaginary coherency provided a frequency-domain skew representation obtained through a separate estimation procedure [7]. Complex coherency was estimated independently in each fixed 50-s window using discrete prolate spheroidal sequence (DPSS) multitaper spectral estimation [20], with time–half-bandwidth product *NW* = 3 and five tapers.

From the multitaper cross-spectral density matrix *S*(*f*),

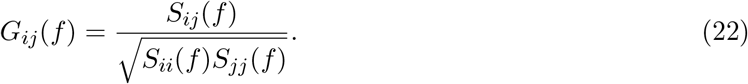

Coherency matrices were averaged over frequency bins within the prespecified band and then across windows. Hermitian symmetry was enforced numerically before extracting the signed imaginary part,

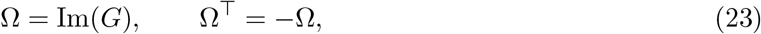

which was factorized as

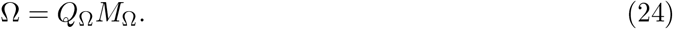

All analyses used the full numerical skew support: 58 dimensions for RestCog (*p* = 59), 58 for ds007176, and 64 for SRM and Dortmund. Raw Ω, *Q*_Ω_, and *M*_Ω_ were evaluated using the matching and fixed alpha–beta fusion rules defined above. The operator *K* contains frequency weighting induced by temporal differencing and time-domain normalization, whereas Ω is estimated directly from normalized signed imaginary spectral interaction.

### 2.8 Finite-lag extension

For lag *τ*,

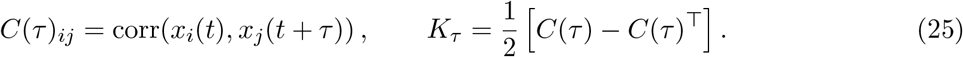

Under second-order stationarity, *C*(−*τ*) = *C*(*τ*)^*T*^, so *K*_*τ*_ = [*C*(*τ*) − *C*(−*τ*)]*/*2 and *K*_*−τ*_ = −*K*_*τ*_.

Discrete lags were *τ*_*ℓ*_ = ℓ*/f*_*s*_, with the horizon fixed a priori as

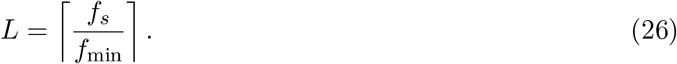

For alpha at *f*_*s*_ = 128 Hz and *f*_min_ = 8 Hz, *L* = 16, corresponding to a 125-ms horizon. No lag was selected from cross-session performance. The lagged operators were assembled into the skew block-Toeplitz matrix

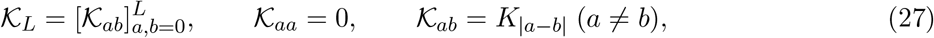

and factorized as K_*L*_ = *Q*_*K*_*M*_*K*_ using the same skew-polar construction.

### 2.9 Mechanistic and specificity controls

Temporal parity was tested using stationary derivative cross-moments

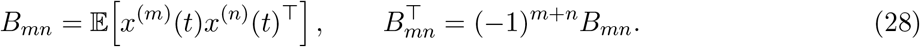

Thus odd total derivative order produces a skew-symmetric cross-moment, whereas even total derivative order produces a symmetric one. In the prespecified RestCog ses01→ses03 analysis, the pairings (0, 1), (0, 2), and (1, 2) were evaluated. The odd-parity operators were projected onto their skew components, whereas the (0, 2) operator was projected onto its symmetric component. A parity-preserving normalization was used, with no derivative-order or rank search.

Three additional controls tested alternative explanations. First, the derivative was recomputed in the Fourier domain, *ℱ*{*ẋ*} (*f*) = *i*2*πf* ℱ{*x*} (*f*). Second, the prespecified *Q*_*T*_ matrices were compared using the spectral operator norm, *d*_op_(*Q*_1_, *Q*_2_) = ∥*Q*_1_ − *Q*_2_ ∥_2_. Third, a one-sided triangular lag embedding was antisymmetrized before polar decomposition to test whether arbitrary skew lifting benefits automatically from normalization. None of these controls introduced performance-based parameter selection.

## 3 Results

### 3.1 Zero-lag reference

Zero-lag Pearson correlation provided a strong symmetric reference (Table 2). Replacing Frobenius distance with the affine-invariant Riemannian metric (AIRM) did not improve verification AUC in any of the five comparisons; CMC@1 improved only in ds007176. The additional Dortmund edge-vector control reached CMC@1 = 0.4709 and AUC = 0.8994. Complete CMC@5 and mean-rank results are reported in Supplementary Table S1.

**Table 2:** Zero-lag Pearson-correlation reference geometry. Cross-session CMC@1 and AUC are shown for Frobenius distance and AIRM, with fixed shrinkage *λ* = 0.01.

| Dataset | Comparison | CMC@1 Frob. | CMC@1 AIRM | AUC Frob. | AUC AIRM |
| --- | --- | --- | --- | --- | --- |
| RestCog | 01–02 | 0.9833 | 0.9833 | 0.9880 | 0.9831 |
| RestCog | 01–03 | 0.4333 | 0.4000 | 0.8864 | 0.7465 |
| ds007176 | V0–V3 | 0.3333 | 0.4074 | 0.7235 | 0.6823 |
| SRM | t1–t2 | 0.7857 | 0.7143 | 0.9565 | 0.9349 |
| Dortmund | 1–2 | 0.4175 | 0.2816 | 0.9050 | 0.8110 |

### 3.2 The empirical state–derivative operator is predominantly time-odd

Although *T* was constructed without imposing antisymmetry, its polar orientation changed only minimally after projection onto the exact skew component *K*. The Frobenius cosine between *Q*_*T*_ and *Q*_*K*_ was 0.999603 in RestCog on the common active support and 0.999252 in ds007176. Antisymmetrization therefore primarily isolated geometry already present in *T*.

A complementary frequency-domain derivative control supported the spectral interpretation. In RestCog, median cos_*F*_ (*K, D*_norm_) = 0.9987. This near-collinearity agrees with the predicted link between the local time-odd operator and frequency-weighted imaginary cross-spectral interaction. Detailed *T*, *Q*_*T*_, and spectral-derivative diagnostics are reported in Supplementary Sections S3–S4.

### 3.3 Polar orientation of the local time-odd operator improves longitudinal discriminability

Table 3 summarizes the principal *K* → *Q*_*K*_ comparison.

**Table 3:**
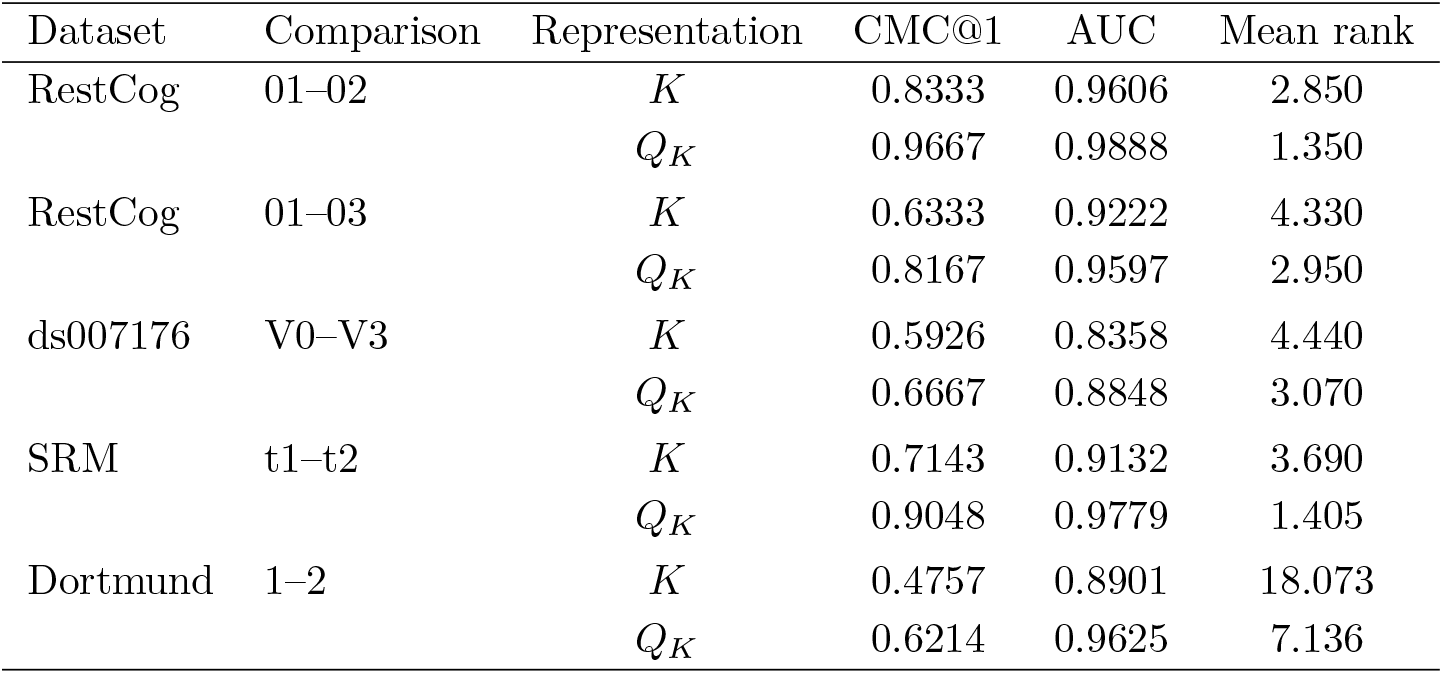
Principal cross-session performance of the raw local time-odd operator *K* and its polar orientation *Q*_*K*_. Raw *K* was matched using strict-upper-triangle Pearson correlation and *Q*_*K*_ using full-matrix Frobenius cosine.

| Dataset | Comparison | Representation | CMC@1 | AUC | Mean rank |
| --- | --- | --- | --- | --- | --- |
| RestCog | 01–02 | $K$ | 0.8333 | 0.9606 | 2.850 |
| | | $Q_K$ | 0.9667 | 0.9888 | 1.350 |
| RestCog | 01–03 | $K$ | 0.6333 | 0.9222 | 4.330 |
| | | $Q_K$ | 0.8167 | 0.9597 | 2.950 |
| ds007176 | V0–V3 | $K$ | 0.5926 | 0.8358 | 4.440 |
| | | $Q_K$ | 0.6667 | 0.8848 | 3.070 |
| SRM | t1–t2 | $K$ | 0.7143 | 0.9132 | 3.690 |
| | | $Q_K$ | 0.9048 | 0.9779 | 1.405 |
| Dortmund | 1–2 | $K$ | 0.4757 | 0.8901 | 18.073 |
| | | $Q_K$ | 0.6214 | 0.9625 | 7.136 |

Polar orientation increased AUC and CMC@1 in every principal local comparison, while mean genuine rank decreased throughout. The effect persisted from the short-interval RestCog comparison to Dortmund, where 206 participants were recorded approximately five years apart: AUC increased from 0.8901 for *K* to 0.9625 for *Q*_*K*_, and CMC@1 from 0.4757 to 0.6214. The same direction was reproduced in the harmonized 4 × 50-s SRM analysis.

The positive factor *M*_*K*_ did not reproduce the orientation advantage (Supplementary Tables S3–S4). Thus stronger cohort-level discrimination was associated with normalized orientation rather than with greater absolute within-subject similarity or the positive magnitude factor.

### 3.4 Frequency-domain replication with imaginary coherency

We next applied the same polar transformation to the spectral skew operator Ω = Im(*G*). In the matched-support RestCog EC alpha analysis, the median same-recording Frobenius cosine was 0.9666 for (*K*, Ω) but only 0.5113 for (*Q*_*K*_, *Q*_Ω_). The two constructions therefore share substantial raw skew structure while yielding non-identical normalized orientations.

Table 4 summarizes the spectral results.

**Table 4:** Cross-session performance of signed imaginary coherency Ω = Im(*G*) and its full-support polar orientation *Q*_Ω_ in alpha and beta. RestCog used the separate EO ses01–ses03 branch; the other cohorts used eyes-closed recordings.

| Dataset | Comparison | Band | CMC@1($\Omega$) | CMC@1($Q_\Omega$) | AUC($\Omega$) | AUC($Q_\Omega$) | $\Delta\text{AUC}$ |
| --- | --- | --- | --- | --- | --- | --- | --- |
| RestCog | 01–03 EO | alpha | 0.3448 | 0.5517 | 0.8530 | 0.8916 | +0.0386 |
|  |  | beta | 0.3103 | 0.4483 | 0.7982 | 0.8743 | +0.0761 |
| ds007176 | V0–V3 CE | alpha | 0.5926 | 0.7407 | 0.8292 | 0.8853 | +0.0561 |
|  |  | beta | 0.2963 | 0.4444 | 0.7119 | 0.8225 | +0.1106 |
| SRM | t1–t2 EC | alpha | 0.6905 | 0.8333 | 0.9106 | 0.9814 | +0.0708 |
|  |  | beta | 0.5238 | 0.6667 | 0.8116 | 0.9324 | +0.1208 |
| Dortmund | 1–2 EC | alpha | 0.4709 | 0.4612 | 0.8977 | 0.9437 | +0.0460 |
|  |  | beta | 0.5049 | 0.5340 | 0.8927 | 0.9444 | +0.0517 |

Across all four datasets, *Q*_Ω_ increased verification AUC relative to Ω in both alpha and beta (8/8 comparisons). CMC@1 increased in 7/8 comparisons; the exception was Dortmund alpha, where AUC nevertheless increased from 0.8977 to 0.9437. The harmonized SRM analysis reproduced the effect in both bands. The positive factor *M*_Ω_ did not reproduce this advantage (Supplementary Table S5), providing an independently estimated frequency-domain instance of the same orientation effect.

### 3.5 Matched-metric robustness control

To exclude matcher choice as an explanation, raw and polar operators were re-evaluated using identical full-matrix Frobenius-cosine and strict-upper-triangle Pearson readouts. Verification AUC increased after polar normalization in all 40 matched raw-to-polar comparisons; EER and mean genuine rank decreased in all 40, while CMC@1 increased in 38 of 40. The only CMC@1 exceptions involved Dortmund alpha-band Ω under the two matched similarity definitions; AUC, EER, CMC@5, and mean rank improved in both cases.

The result was consistent across operator families, datasets, bands, fusion, and similarity definitions. For example, in RestCog 01–03 EC, Frobenius-cosine AUC increased from 0.9214 to 0.9597 for *K* → *Q*_*K*_ and from 0.9232 to 0.9692 for Ω → *Q*_Ω_. In the harmonized SRM fusion control, AUC increased from 0.9273 to 0.9864 for *K* → *Q*_*K*_ and from 0.9242 to 0.9910 for Ω → *Q*_Ω_. In Dortmund fusion, the corresponding Frobenius-cosine gains were 0.9417 to 0.9855 and 0.9460 to 0.9766. Strict-upper-triangle Pearson produced the same direction of effect. All raw–polar pairs within these prespecified controls shared identical temporal support. Selected endpoints spanning the cross-dataset control are reported in Supplementary Table S6.

### 3.6 Cross-band complementarity of polar orientation

Fixed equal-weight alpha–beta fusion tested whether the two bands contained complementary orientation information. Table 5 summarizes the local *Q*_*K*_ results.

**Table 5:**
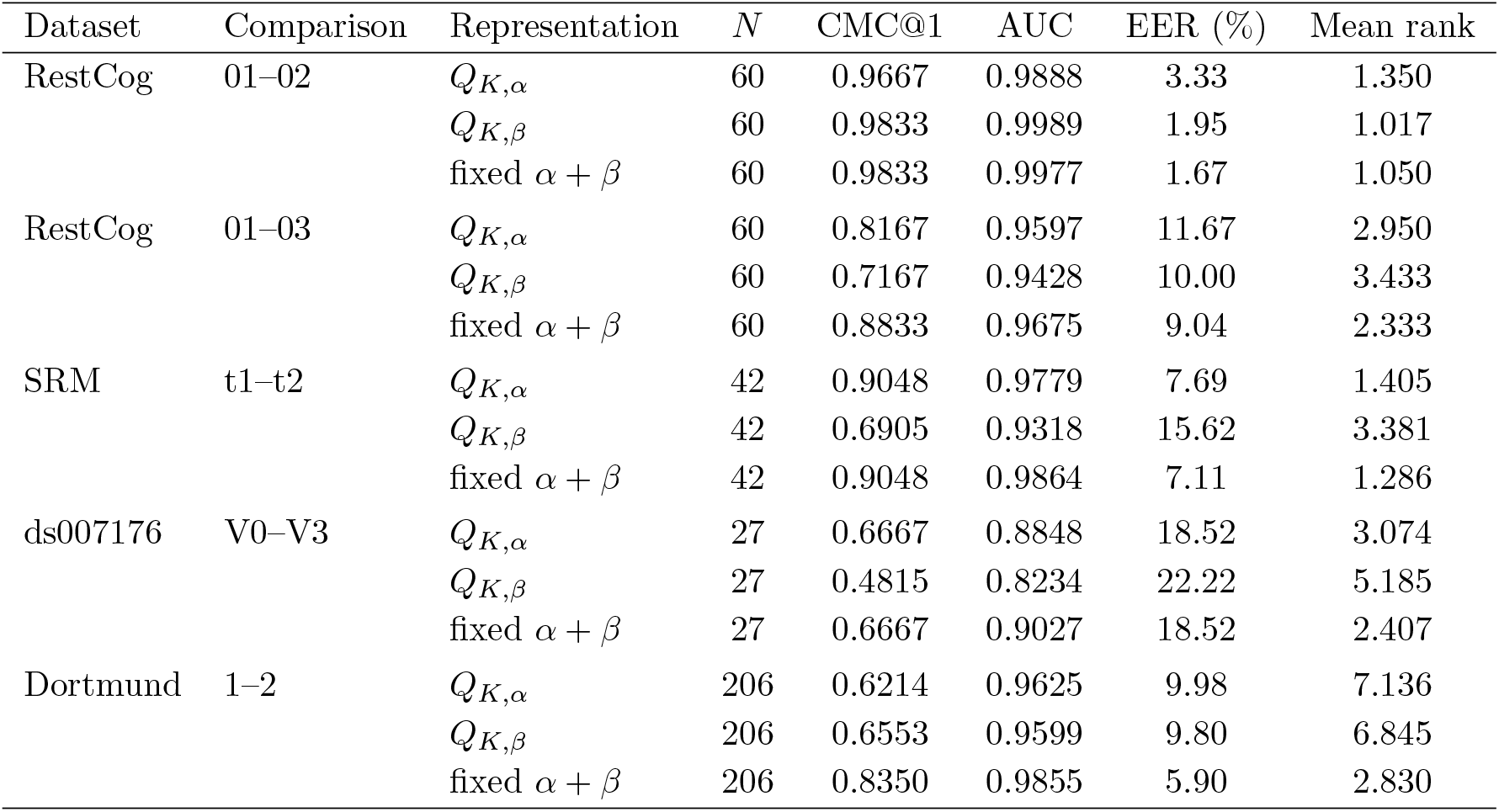
Cross-session performance of full-support *Q*_*K*_ in alpha, beta, and fixed equal-weight alpha– beta score fusion.

Fusion was most beneficial in the more demanding longitudinal comparisons. In RestCog 01–03, CMC@1 increased from 0.8167 in alpha to 0.8833 after fusion. In Dortmund, fusion increased CMC@1 from 0.6214 and 0.6553 in the individual bands to 0.8350, with AUC = 0.9855 and EER = 5.90%. In SRM and ds007176, fusion improved AUC and mean rank without increasing CMC@1 relative to alpha.

Figure 2 shows that the Dortmund fusion gain corresponds to a larger angular margin between genuine matches and their nearest impostors.

The spectral orientation *Q*_Ω_ showed a corresponding cross-band pattern (Table 6).

**Table 6:** Cross-band performance of full-support *Q*_Ω_. ΔAUC is fusion minus the better single-band AUC; CMC@1 and mean rank refer to the fused representation.

| Dataset / comparison | AUC $\alpha$ | AUC $\beta$ | AUC $\alpha + \beta$ | $\Delta\text{AUC}$ | CMC@1 | Mean rank |
| --- | --- | --- | --- | --- | --- | --- |
| RestCog 01–03 EO | 0.8916 | 0.8743 | 0.9304 | +0.0388 | 0.6207 | 3.983 |
| ds007176 V0–V3 CE | 0.8853 | 0.8225 | 0.9095 | +0.0242 | 0.6296 | 2.444 |
| SRM t1–t2 EC | 0.9814 | 0.9324 | 0.9910 | +0.0096 | 0.9286 | 1.119 |
| Dortmund 1–2 EC | 0.9437 | 0.9444 | 0.9766 | +0.0322 | 0.7524 | 4.240 |

Fixed fusion increased *Q*_Ω_ AUC in all four datasets. In the separate RestCog EO branch, the fusion gain over alpha was supported by paired inference (Holm-adjusted *p* = 0.0206). Cross-band complementarity was therefore reproducible but dataset-dependent and remained secondary to the within-band raw-to-polar effect.

### 3.7 Finite-lag and mechanistic controls

The finite-lag extension reproduced the polar effect in every evaluated condition. AUC increased from 0.9673 to 0.9904 in RestCog 01–02, from 0.9371 to 0.9631 in RestCog 01–03, from 0.7876 to 0.8898 in ds007176, and from 0.9276 to 0.9822 in SRM. Its incremental benefit over the compact local orientation was modest. Complete identification results are reported in Supplementary Table S7.

Derivative parity provided the principal mechanistic specificity control. The odd-parity (0, 1) and (1, 2) operators had raw AUCs of 0.9222 and 0.9221 and polar AUCs of 0.9598 and 0.9664, respectively; CMC@1 increased from 0.6333 to 0.8167 for both. By contrast, the even-parity (0, 2) polar factor was non-discriminative, with AUC = 0.5000. Thus the polar effect was present for odd temporal parity, disappeared when the skew time-odd geometry was removed, and reappeared when odd parity was restored. Additional Fourier-derivative, operator-norm, positive-factor, and antisymmetrized triangular-lag controls are reported in Supplementary Sections S3–S4 and S7.

## 4 Discussion

### 4.1 Polar orientation of local time-odd dynamics

The principal finding is that polar normalization improves the longitudinal discriminability of the local time-odd operator. In the factorization *K* = *Q*_*K*_*M*_*K*_, each active canonical skew plane is weighted in *K* by its singular magnitude *σ*_*j*_, whereas *Q*_*K*_ replaces *σ*_*j*_*J* by *J*. Polar normalization therefore removes skew-plane magnitude weighting while retaining normalized orientation. Across the longitudinal comparisons, *Q*_*K*_ outperformed the raw operator *K*, whereas the positive factor *M*_*K*_ did not reproduce the same advantage. A plausible interpretation is that session-sensitive magnitude weighting is partially removed while distributed orientation geometry is retained. This does not imply that magnitude is intrinsically uninformative; rather, it did not carry the same longitudinal advantage in the present comparisons.

Improved discrimination should not be interpreted as smaller absolute cross-session change. As illustrated in Figure 1, the representative *Q*_*K*_ pair was less aligned across sessions than the corresponding *K* and *M*_*K*_ pairs, yet was more discriminative at the cohort level. Polar normalization therefore reorganizes relative genuine–impostor geometry rather than simply contracting within-subject distances.

**Figure 1:**
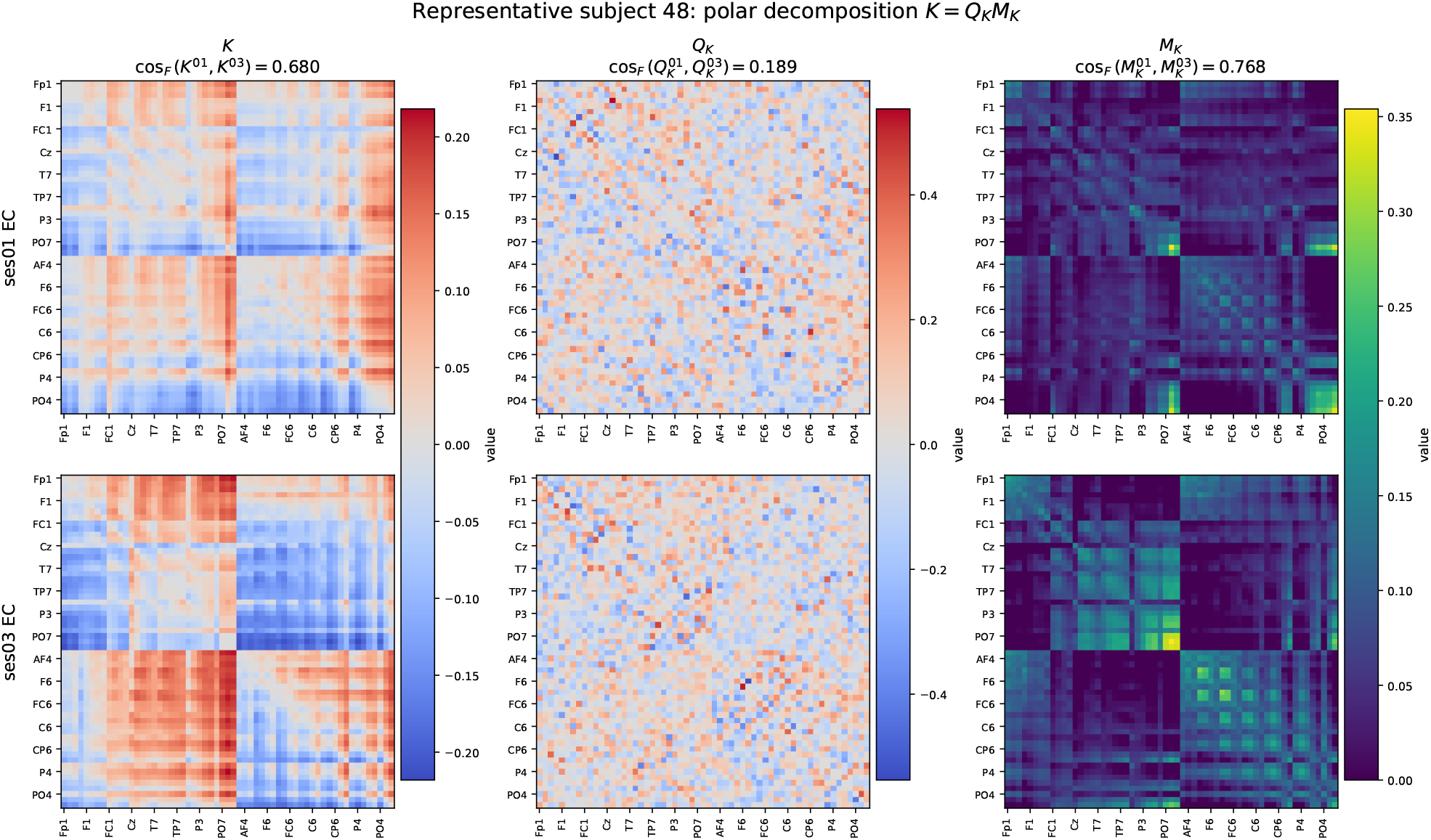
Representative RestCog ses01–ses03 decomposition *K* = *Q*_*K*_*M*_*K*_. The participant was selected because its genuine cross-session *Q*_*K*_ Frobenius cosine was closest to the cohort median. For this participant, cos_*F*_ (*K*^01^, *K*^03^) = 0.680, 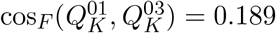, and 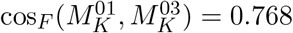. The polar gain therefore reflects relative genuine–impostor geometry rather than greater absolute within-subject similarity.

**Figure 2:**
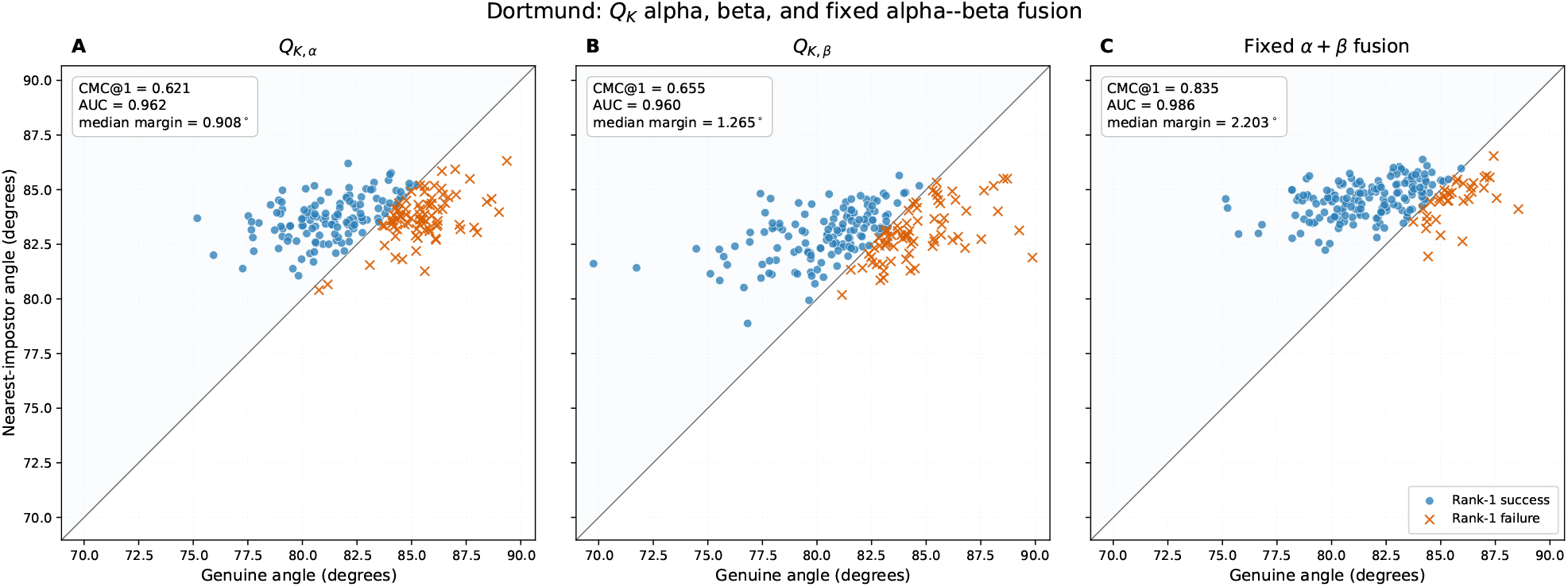
Dortmund nearest-neighbor angular geometry for *Q*_*K*_ in alpha, beta, and fixed alpha–beta fusion. Points above the diagonal indicate correct rank-1 matches. The median genuine-to-nearest-impostor margin increased from 0.908^*°*^ in alpha and 1.265^*°*^ in beta to 2.203^*°*^ after fusion, while CMC@1 increased to 0.835.

The orientation effect was also not created by explicit antisymmetrization. The empirical state– derivative operator *T* was already strongly dominated by its skew component, and *Q*_*T*_ was nearly identical to *Q*_*K*_. Together with the near-collinearity between *K* and the normalized spectral-derivative control, this supports viewing *K* as a time-domain realization of frequency-weighted imaginary cross-spectral interaction rather than as an arbitrary algebraic projection.

Polar normalization is not guaranteed to improve longitudinal matching. When an active skew singular value approaches zero, its orientation becomes poorly conditioned and can be sensitive to perturbation. The observed benefit is therefore an empirical property of the EEG operators studied here rather than a mathematical consequence of the polar map itself.

### 4.2 Spectral replication and representation-level robustness

The imaginary-coherency operator Ω = Im(*G*) provides a frequency-domain realization of skew EEG structure obtained through a separate estimation procedure [7]. It is analytically linked to *K* through the imaginary cross-spectrum but differs in frequency weighting, normalization, and estimation. The strong same-recording alignment of the raw operators, together with the substantially lower alignment of their polar factors, is consistent with related but non-equivalent representations.

Despite these differences, Ω → *Q*_Ω_ reproduced the same raw-to-polar AUC improvement as *K* → *Q*_*K*_ in both alpha and beta across all four spectral datasets. This extends the effect beyond one particular estimator and supports the interpretation that normalized skew orientation, rather than the numerical details of the local state–derivative construction, carries the relevant longitudinal information.

The matched-metric controls provide a particularly important representation-level test. When the same similarity definition was applied before and after normalization, verification AUC increased and both EER and mean genuine rank decreased in all 40 raw-to-polar comparisons; CMC@1 increased in 38 of 40. Raw and polar representations shared identical temporal support within these dedicated controls. The effect therefore cannot be attributed to comparing raw operators with Pearson correlation and polar factors with Frobenius cosine.

The zero-lag reference provides a complementary negative control. Replacing Frobenius comparison of Pearson-correlation matrices with the affine-invariant Riemannian metric did not improve verification AUC in any of the five cross-session comparisons. Thus the principal gain is not explained simply by changing the metric on an otherwise unchanged representation; the time-odd polar construction changes the represented information itself.

### 4.3 Temporal parity and finite-lag interpretation

Derivative parity provides a mechanistic specificity test. Under stationarity, odd total derivative order produces a skew-symmetric cross-moment, whereas even total derivative order produces a symmetric one. Consistent with this prediction, the odd-parity (0, 1) and (1, 2) constructions showed strong polar discriminability, whereas the even-parity (0, 2) polar factor was non-discriminative. In this prespecified control, the effect tracked temporal parity rather than differentiation or orthogonalization in general.

The finite-lag analysis places the local state–derivative operator within a broader time-odd correlation geometry. Polar normalization improved the finite-lag representation in every evaluated condition, but its incremental benefit over the compact *p* × *p* local orientation was modest. The larger embedding therefore serves mainly as an extension test: the orientation principle persists beyond the local limit without requiring a substantially more complex temporal representation.

### 4.4 Cross-band complementarity

Alpha- and beta-band polar orientations were not fully redundant. Fixed equal-weight fusion provided complementary longitudinal information in several cohorts, with the clearest identification gain in Dortmund. For recordings separated by approximately five years, the fused *Q*_*K*_ reached CMC@1 = 0.8350, AUC = 0.9855, and EER = 5.90%. The nearest-neighbor analysis showed that this gain reflected a larger margin between genuine matches and their closest impostors.

The effect was nevertheless dataset-dependent. In SRM and ds007176, local fusion improved AUC and mean rank without increasing CMC@1 relative to alpha, whereas the spectral *Q*_Ω_ analysis showed fusion-related AUC gains across all four datasets. Cross-band complementarity should therefore be viewed as a secondary extension of the main result: alpha and beta can provide partly distinct orientation information, but fusion is neither necessary for the raw-to-polar advantage nor guaranteed to improve every metric or cohort.

### 4.5 Longitudinal interpretation and limitations

The two RestCog comparisons suggest that polar orientation may be particularly useful when cross-session variability is substantial. Zero-lag correlation was near ceiling in the short-interval comparison but degraded in the more demanding ses01–ses03 comparison, whereas the time-odd polar representation remained strongly discriminative. This contrast is descriptive and should not be interpreted as a temporal decay law.

Dortmund provides the strongest long-interval stress test. In 206 participants with recordings separated by approximately five years, AUC increased from 0.8901 for *K* to 0.9625 for *Q*_*K*_. This does not imply invariance over five years; rather, it shows that substantial subject-specific organization remains recoverable across a much longer interval. The result is consistent with previous evidence for longitudinal EEG permanence [1–3], while extending that literature to a time-odd representation that explicitly separates interaction magnitude from orientation.

Cross-session matching is used here as a stringent assay of longitudinal reproducibility, not as evidence for a neural “identity” invariant. All analyses were performed in sensor space, where head anatomy, volume conduction, electrode placement, reference choice, and other technical or partly non-neural factors may contribute to subject-specific structure. Signed imaginary coherency reduces sensitivity to instantaneous zero-phase coupling but does not eliminate these confounds [7, 21]. Likewise, the sign of a skew interaction should not be interpreted as causal, synaptic, or as evidence of directed physiological influence.

The representation is not invariant to arbitrary montage or reference transformations, and its behavior under substantial changes in sensor configuration remains to be established. Whether the same orientation geometry persists after source reconstruction or across different sensor layouts is an important question for future work. Polar orientation can also become unstable when active skew singular values approach zero; full-support factorization avoids rank optimization but not the conditioning limits of the polar map.

The datasets differ in cohort size, hardware, montage, protocol, recording duration, and inter-session interval, so between-dataset performance differences cannot be attributed to elapsed time alone. Frequency analysis was restricted to alpha and beta, and fusion was deliberately fixed rather than optimized. The smaller ds007176 cohort provides an independent replication but carries less statistical weight than Dortmund. Verification scores are statistically dependent because many genuine and impostor comparisons share participants; pairwise scores should therefore not be treated as independent observations. Subject-level identification and paired analyses provide complementary evidence, while EER comparisons remain descriptive unless uncertainty is estimated using a dependence-preserving procedure. More generally, the repeated direction of the raw-to-polar effect across frozen cohorts and controls is emphasized here over a claim of universal statistical superiority.

## 5 Conclusion

This study identifies polar orientation as a reproducible representation of longitudinal subject-specific structure in multichannel time-odd EEG dynamics. The local state–derivative operator retained the near-skew geometry predicted under stationarity, and its exact skew component *K* = (*T* − *T*^*T*^)*/*2 therefore isolated structure already present in the data. Polar factorization separated skew-plane magnitude weighting from orientation, and the orientation factor *Q*_*K*_ consistently provided stronger cross-session discriminability than the raw operator across four longitudinal EEG datasets, including the Dortmund cohort with recordings separated by approximately five years.

The same raw-to-polar effect was independently reproduced for the frequency-domain skew operator Ω = Im(*G*) in both alpha and beta. It also persisted when raw and polar representations were evaluated with identical similarity definitions, with verification AUC improving in all 40 matched-metric comparisons. Derivative-parity and finite-lag controls further linked the effect to odd temporal structure and showed that the orientation principle extends beyond the local state–derivative limit. Fixed alpha–beta fusion provided additional complementary information in some cohorts, but was secondary to the within-band raw-to-polar effect.

Together, these findings support a geometric distinction between the magnitude and orientation of multichannel time-odd EEG interactions. Removing interaction-magnitude weighting does not imply that magnitude is uninformative; rather, in the present longitudinal comparisons, normalized orientation provided the more discriminative cross-session representation. The results therefore motivate polar geometry as a compact framework for studying reproducible temporal organization in multichannel EEG.

