## Supplementary Information for "Polar Geometry of Time-Odd EEG Dynamics"

Dmitry Goldstein

August 31, 2026

#### S1 Scope and analysis conventions

This Supplementary Information provides the numerical reference results and specificity controls cited in the main manuscript. All reported analyses use prespecified enrollment–probe protocols. No classifier, metric learning, identity-specific template fitting, score calibration, fusion-weight tuning, rank scan, or probe-dependent selection is introduced here.

The notation follows the main manuscript. The local empirical state–derivative operator is denoted by $T$, its exact time-odd projection by $K = (T - T^\top)/2$, and the corresponding polar decompositions by $T = Q_T M_T$ and $K = Q_K M_K$. The frequency-domain skew operator is $\Omega = \mathrm{Im}(G)$, with $\Omega = Q_\Omega M_\Omega$. For matched-metric controls, the same similarity definition is applied before and after polar normalization. Fixed alpha–beta fusion is always

$$\Gamma_{\alpha\beta} = \tfrac{1}{2}\Gamma_\alpha + \tfrac{1}{2}\Gamma_\beta, \tag{S1}$$

with no score normalization or learned weighting.

#### S2 Zero-lag reference geometry

Table S1 reports the complete zero-lag identification and ranking statistics used as the conventional symmetric reference. Frobenius distance is computed directly on Pearson correlation matrices. The affine-invariant Riemannian metric (AIRM) uses fixed shrinkage $\lambda = 0.01$. The Dortmund edge control uses Pearson correlation of vectorized correlation-matrix edges.

**Table S1:** Complete zero-lag reference results. Higher values are better for CMC@1, CMC@5, and AUC; lower mean rank is better.

| Dataset | Comparison | Readout | CMC@1 | CMC@5 | AUC | Mean rank |
|---|---|---|---|---|---|---|
| RestCog | 01–02 | Frobenius | 0.9833 | 1.0000 | 0.9880 | 1.05 |
| | | AIRM | 0.9833 | 0.9833 | 0.9831 | 1.15 |
| RestCog | 01–03 | Frobenius | 0.4333 | 0.8000 | 0.8864 | 4.97 |
| | | AIRM | 0.4000 | 0.6000 | 0.7465 | 8.88 |
| ds007176 | V0–V3 | Frobenius | 0.3333 | 0.5185 | 0.7235 | 8.11 |
| | | AIRM | 0.4074 | 0.6296 | 0.6823 | 8.56 |
| SRM | t1–t2 | Frobenius | 0.7857 | 0.9524 | 0.9565 | 1.69 |
| | | AIRM | 0.7143 | 0.8571 | 0.9349 | 3.19 |
| Dortmund | 1–2 | Edge Pearson | 0.4709 | 0.6165 | 0.8994 | 16.85 |
| | | Frobenius | 0.4175 | 0.6699 | 0.9050 | 15.22 |
| | | AIRM | 0.2816 | 0.4223 | 0.8110 | 30.43 |

AIRM did not improve verification AUC over Frobenius in any of the five cross-session comparisons. In Dortmund, the additional edge-vector control was in the same broad performance range as the full-matrix zero-lag reference.

### S3 Local operator and polar-factor controls

#### S3.1 Near-skew structure and the role of antisymmetrization

The empirical directed operator was already almost entirely time-odd. In RestCog, the skew-energy fraction of the independently differentiated operator was 0.999862, and the corresponding polar factor was essentially unchanged. The Frobenius cosine between the polar factors of $T$ and $K$ was

$$\cos_F(Q_T, Q_K) = 0.999603 \tag{S2}$$

in RestCog after restriction to the common active support and

$$\cos_F(Q_T, Q_K) = 0.999252 \tag{S3}$$

in ds007176. These values show that explicit antisymmetrization isolates an orientation already present in the empirical directed operator rather than creating a new orientation geometry.

A complementary exact-skew normalization control in RestCog produced the results in Table S2. Numerically symmetrizing the correlation construction before forming $K$ left both the raw skew operator and its polar orientation essentially unchanged.

**Table S2:** RestCog 01–03 exact-skew normalization control. The two skew constructions are numerically equivalent at the reported precision.

| Representation | CMC@1 | CMC@5 | AUC | Mean rank |
|---|---|---|---|---|
| $K_{\text{corr}}$ | 0.6333 | 0.8000 | 0.9222 | 4.333 |
| $Q_{K_{\text{corr}}}$ | 0.8167 | 0.8833 | 0.9597 | 2.950 |
| $M_{K_{\text{corr}}}$ | 0.6667 | 0.8667 | 0.9039 | 4.483 |
| $K_{\text{sym}}$ | 0.6333 | 0.8000 | 0.9222 | 4.333 |
| $Q_{K_{\text{sym}}}$ | 0.8167 | 0.8833 | 0.9598 | 2.950 |
| $M_{K_{\text{sym}}}$ | 0.6667 | 0.8667 | 0.9039 | 4.483 |
| $Q_T$ reference | 0.8167 | 0.8833 | 0.9590 | 2.967 |

For the two exact-skew implementations,

$$\operatorname{median} \cos_F(K_{\text{corr}}, K_{\text{sym}}) = 1.000000, \tag{S4}$$
$$\operatorname{median} \cos_F(Q_{K_{\text{corr}}}, Q_{K_{\text{sym}}}) = 0.999996. \tag{S5}$$

#### S3.2 Positive polar factors do not reproduce the orientation gain

The positive factor was less discriminative than the polar orientation in the completed local magnitude controls. Table S3 reports verification AUC for the three longitudinal comparisons with complete, protocol-matched $M_T/M_K$ exports.

For Dortmund, the full local decomposition ledger is shown in Table S4. The two orientation factors form a high-performance tier, whereas the corresponding positive factors remain close to or below the raw operators.

**Table S3:** Local positive-factor controls. The positive factors do not reproduce the verification performance of $Q_K$.

| Dataset | Comparison | AUC($M_T$) | AUC($M_K$) | AUC($Q_K$) |
|---|---|---|---|---|
| RestCog | 01–03 | 0.9045 | 0.9039 | 0.9597 |
| ds007176 | V0–V3 | 0.7773 | 0.7776 | 0.8848 |
| Dortmund | 1–2 | 0.8602 | 0.8594 | 0.9625 |

**Table S4:** Dortmund prespecified local-operator decomposition, $N = 206$.

| Representation | CMC@1 | CMC@5 | AUC | Mean rank |
|---|---|---|---|---|
| $T$ | 0.4612 | 0.6214 | 0.8871 | 18.59 |
| $Q_T$ | 0.6117 | 0.7621 | 0.9625 | 7.17 |
| $M_T$ | 0.4951 | 0.6650 | 0.8602 | 21.64 |
| $K$ | 0.4757 | 0.6262 | 0.8901 | 18.07 |
| $Q_K$ | 0.6214 | 0.7573 | 0.9625 | 7.14 |
| $M_K$ | 0.5000 | 0.6602 | 0.8594 | 21.78 |

#### S4 Frequency-domain magnitude and mechanistic controls

The spectral branch uses the signed skew operator $\Omega = \mathrm{Im}(G)$. Table S5 collects the complete magnitude-factor metrics available on prespecified branches whose temporal support matches the corresponding principal result. Within these available matched-support controls, the orientation factor remains substantially above $M_\Omega$ in verification AUC.

**Table S5:** Frequency-domain positive-factor controls on prespecified matched-support branches. "Fusion" is the fixed 0.5/0.5 alpha–beta score average.

| Dataset | Band | Representation | CMC@1 | CMC@5 | AUC | EER (%) | Mean rank |
|---|---|---|---|---|---|---|---|
| RestCog EO | alpha | $M_\Omega$ | 0.3448 | 0.5000 | 0.7689 | – | 11.086 |
| | beta | $M_\Omega$ | 0.2759 | 0.4483 | 0.6785 | – | 14.259 |
| ds007176 | alpha | $M_\Omega$ | 0.4815 | 0.7037 | 0.7761 | 30.34 | 5.185 |
| | beta | $M_\Omega$ | 0.2593 | 0.5185 | 0.7029 | 36.32 | 8.222 |
| | fusion | $M_\Omega$ | 0.4074 | 0.7037 | 0.7732 | 29.20 | 5.741 |

The local and spectral skew operators are mathematically linked through the imaginary cross-spectrum but are not equivalent estimators. In the matched-support RestCog alpha analysis, the same-recording raw operators were strongly aligned,

$$\operatorname{median} \cos_F(K, \Omega) = 0.9666, \tag{S6}$$

whereas their normalized orientation factors were much less aligned,

$$\operatorname{median} \cos_F(Q_K, Q_\Omega) = 0.5113. \tag{S7}$$

Thus the two constructions share substantial raw time-odd structure while their polar normalization exposes partially distinct orientation geometry.

A separate spectral-derivative construction provides a direct mechanistic check on the local state–derivative interpretation. The normalized frequency-weighted imaginary cross-spectral operator was almost collinear with the time-domain $K$ in RestCog,

$$\operatorname{median} \cos_F(K, D_{\mathrm{norm}}) = 0.9987. \tag{S8}$$

This supports the expected spectral identity without requiring the two estimators to be numerically identical in their final normalized forms.

#### S5 Matched-metric robustness control

The matched-metric analysis applies exactly the same readout to each raw skew operator and its polar orientation: either normalized full-matrix Frobenius cosine or Pearson correlation of the signed strict upper triangle. Across the complete prespecified control, verification AUC increased after polar normalization in all 40 raw-to-polar comparisons; EER and mean genuine rank decreased in all 40, while CMC@1 increased in 38 of 40. The only CMC@1 exceptions involved Dortmund alpha-band $\Omega$ under the two matched similarity definitions; AUC, EER, CMC@5, and mean rank improved in both cases.

Table S6 reports selected verified endpoints spanning all four datasets, both operator families, and both similarity definitions, including the harmonized SRM $4 \times 50$-s control. The directional summary above refers to the complete 40-comparison prespecified analysis; the table is intentionally limited to representative cross-dataset endpoints rather than reproducing the full score ledger.

**Table S6:** Selected matched-metric robustness endpoints used to anchor the cross-dataset summary. Raw and polar representations within each row use the same matcher and temporal support.

| Dataset | Band | Family | Matcher | AUC raw | AUC polar | ΔAUC |
|---|---|---|---|---|---|---|
| RestCog 01–03 EC | alpha | $K \to Q_K$ | Frobenius | 0.9214 | 0.9597 | +0.0383 |
| | alpha | $K \to Q_K$ | Pearson | 0.9222 | 0.9583 | +0.0361 |
| | alpha | $\Omega \to Q_\Omega$ | Frobenius | 0.9232 | 0.9692 | +0.0460 |
| | alpha | $\Omega \to Q_\Omega$ | Pearson | 0.9250 | 0.9667 | +0.0417 |
| SRM t1–t2 EC, $4 \times 50$ s | fusion | $K \to Q_K$ | Frobenius | 0.9273 | 0.9864 | +0.0591 |
| | fusion | $K \to Q_K$ | Pearson | 0.9267 | 0.9869 | +0.0602 |
| | fusion | $\Omega \to Q_\Omega$ | Frobenius | 0.9242 | 0.9910 | +0.0668 |
| | fusion | $\Omega \to Q_\Omega$ | Pearson | 0.9233 | 0.9912 | +0.0679 |
| ds007176 V0–V3 | fusion | $K \to Q_K$ | Frobenius | 0.7289 | 0.9027 | +0.1738 |
| | fusion | $K \to Q_K$ | Pearson | 0.8085 | 0.9032 | +0.0947 |
| | fusion | $\Omega \to Q_\Omega$ | Frobenius | 0.7259 | 0.9095 | +0.1836 |
| | fusion | $\Omega \to Q_\Omega$ | Pearson | 0.8028 | 0.9130 | +0.1102 |
| Dortmund 1–2 | fusion | $K \to Q_K$ | Frobenius | 0.9417 | 0.9855 | +0.0438 |
| | fusion | $K \to Q_K$ | Pearson | 0.9433 | 0.9855 | +0.0422 |
| | fusion | $\Omega \to Q_\Omega$ | Frobenius | 0.9460 | 0.9766 | +0.0306 |
| | fusion | $\Omega \to Q_\Omega$ | Pearson | 0.9464 | 0.9767 | +0.0303 |

For Dortmund alpha $\Omega$, the two CMC@1 exceptions were $0.4806 \to 0.4612$ under Frobenius cosine and $0.4709 \to 0.4660$ under strict-upper-triangle Pearson correlation. In both comparisons, verification AUC increased and the remaining ranking and error summaries improved.

#### S6 Finite-lag extension

The finite-lag construction tests whether the same time-odd polar principle persists beyond the local derivative limit. Table S7 reports the complete prespecified identification and verification results. In every evaluated condition, polar normalization increased AUC and decreased mean rank.

The corresponding odd-energy diagnostics are reported in Table S8. The increasing odd fraction with lag in the RestCog and SRM analyses is consistent with the finite-lag operator capturing progressively stronger time-odd structure over the sampled lag horizon.

For SRM, the lifted dimension was 1088 and all evaluated lifted operators were full rank. The maximum relative reconstruction error, maximum $\|Q^\top Q - I\|$ error, and maximum $\|Q^2 + I\|$ error were $2.32 \times 10^{-15}$, $3.52 \times 10^{-15}$, and $3.52 \times 10^{-15}$, respectively.

**Table S7:** Complete finite-lag identification and verification results.

| Dataset | Comparison | Representation | CMC@1 | CMC@5 | AUC | Mean rank |
|---|---|---|---|---|---|---|
| RestCog | 01–02 | $K_L$ | 0.8333 | 0.9000 | 0.9673 | 2.567 |
| | | $Q_{K_L}$ | 0.9667 | 0.9833 | 0.9904 | 1.283 |
| RestCog | 01–03 | $K_L$ | 0.6500 | 0.8667 | 0.9371 | 3.350 |
| | | $Q_{K_L}$ | 0.8333 | 0.9000 | 0.9631 | 2.700 |
| ds007176 | V0–V3 | $K_L$ | 0.6667 | 0.7778 | 0.7876 | 4.519 |
| | | $Q_{K_L}$ | 0.6667 | 0.8519 | 0.8898 | 2.889 |
| SRM | t1–t2 | $K_L$ | 0.7619 | 0.8810 | 0.9276 | 3.190 |
| | | $Q_{K_L}$ | 0.9286 | 1.0000 | 0.9822 | 1.143 |

**Table S8:** Odd-fraction diagnostics for the finite-lag branch.

| Regime | Mean odd enroll. | Mean odd probe | Median lag-1 odd | Median lag-$L$ odd |
|---|---|---|---|---|
| RestCog 01–02 | 0.3292 | 0.3428 | 0.1157 | 0.5051 |
| RestCog 01–03 | 0.3292 | 0.3212 | 0.1083 | 0.4752 |
| ds007176 V0–V3 | 0.3345 | 0.2877 | 0.1025 | – |
| SRM t1–t2 | 0.3215 | 0.3049 | 0.1022 | 0.5018 |

### S7 Mechanistic specificity controls

#### S7.1 Derivative parity

For stationary derivative cross-moments, odd total derivative order is skew-symmetric, whereas even total derivative order is symmetric. The empirical parity experiment follows this prediction (Table S9). The polar effect is present for the odd-parity $(0, 1)$ and $(1, 2)$ constructions, disappears for even parity $(0, 2)$, and reappears when odd parity is restored.

**Table S9:** Derivative-parity control in RestCog.

| Derivative orders | Parity | Representation | Geometry | CMC@1 | AUC |
|---|---|---|---|---|---|
| $(0, 1)$ | odd | raw | time-odd | 0.6333 | 0.9222 |
| | | polar | orientation | 0.8167 | 0.9598 |
| $(0, 2)$ | even | polar | symmetric-limit control | – | 0.5000 |
| $(1, 2)$ | odd | raw | time-odd | 0.6333 | 0.9221 |
| | | polar | orientation | 0.8167 | 0.9664 |

#### S7.2 Negative geometric controls

Table S10 collects two controls that delimit the mechanism. First, replacing the distributed Frobenius readout of $Q_T$ with the spectral operator norm largely destroys discrimination, arguing against a single worst-direction explanation. Second, polar normalization does not automatically improve an arbitrary antisymmetrized lifted matrix: the one-sided triangular-lag construction degrades in the more demanding RestCog 01–03 comparison.

These controls constrain the interpretation of the main effect. Polar normalization is not sufficient by itself; the gain depends on the particular distributed orientation geometry carried by the empirically observed time-odd operator.

**Table S10:** Specificity controls against generic orthogonal normalization or arbitrary antisymmetric lifting.

| Control | Representation / readout | CMC@1 | AUC | Mean rank |
|---|---|---|---|---|
| RestCog 01–02 local orientation | $Q_T$, Frobenius cosine | 0.9667 | 0.9888 | 1.400 |
| RestCog 01–02 operator-norm control | $Q_T$, spectral operator norm | 0.0833 | 0.5912 | 30.30 |
| RestCog 01–03 triangular lag | raw antisymmetrized lift | 0.7833 | 0.9644 | 1.883 |
| RestCog 01–03 triangular lag | polar factor of lift | 0.7500 | 0.9128 | 3.100 |

#### S8 Reproducibility and scope notes

The manuscript combines branches with different recording supports only when they address different, explicitly identified questions. The principal frequency-domain replication uses the prespecified support for each cohort, whereas the matched-metric analysis uses dedicated branches in which raw and polar operators share exactly the same support. For SRM, all current matched-metric results use the harmonized central 200-s support divided into four non-overlapping 50-s windows. No superseded SRM matched-support values are included in the current tables or summary statements.

The main inference likewise does not rely on exploratory rank-truncated optima. Reported polar factors use full numerical support, and cross-band fusion is fixed before evaluation. This Supplementary Information is intended as a numerical audit of the manuscript's prespecified claims rather than as a catalogue of exploratory constructions considered during method development.
